# LAVR-289, a new acyclo-nucleoside phosphonate having broad-spectrum activity against herpesviruses

**DOI:** 10.1101/2025.02.20.639249

**Authors:** Sandrine Kappler-Gratias, Charlotte Quentin-Froignant, Elie Marcheteau, José Fernandez, David Boutolleau, Virginie Garcia, Vincent Roy, Luigi A. Agrofoglio, Franck Gallardo

**Affiliations:** NeoVirTech SAS, Toulouse, France; Sorbonne Université, INSERM U1136, Pierre Louis Institute of Epidemiology and Public Health (IPLESP), THERAVIR team, Paris, France; AP-HP. Sorbonne Université, National Reference Center for Herpesviruses (associated laboratory), Virology Department, Pitié-Salpêtrière Hospital, Paris, France; Unité Mixte de Recherche Institut National de la Santé et de la Recherche Médicale (INSERM) 1037, Centre National de la Recherche Scientifique (CNRS) 5071, Université Toulouse III - Paul Sabatier, Centre de Recherches en Cancérologie de Toulouse (CRCT), Toulouse, France; Institute of Organic and Analytical Chemistry (ICOA UMR 7311), University of Orleans, CNRS, F-45067 Orléans, France

**Keywords:** nucleoside, broad-spectrum antiviral, herpesvirus

## Abstract

Human herpesviruses are latent opportunistic large dsDNA viruses that can have deleterious effect in immunocompromised patients by triggering life-threatening infections. Over the past years, different antivirals have been developed against a variety of herpesviruses, including Acyclovir for herpes simplex viruses (HSV1 and 2) and ganciclovir and letermovir for human cytomegalovirus (hCMV). However, broad-spectrum inhibitors of herpesvirus infections are still missing. Here we report the efficacy of LAVR-289, a new acyclic nucleoside analog, on a broad variety of herpesviruses from human and animal origin. LAVR-289 displays nanomolar efficiency *in vitro*, is active on viral strains resistant to gold standard antivirals and *ex vivo* on reconstituted human skin infected with HSV1. Combined with its anti-poxvirus and anti-adenovirus activity, LAVR-289 can become the next gold standard for the management of opportunistic virus infections in immunocompromised patient.

## INTRODUCTION

DNA viruses belong to several families that can lead to acute life-threatening infection such as poxviruses to life-long infection such as herpesviruses. The vast majority of DNA viruses are composed of double stranded DNA (dsDNA), either in the linear or circular form. DNA viruses hijack cell machinery to replicate and whereas large viruses encode their own polymerase, smallest viruses need host cell support for replication. Among dsDNA virus families infecting mammalian cells, *Orthoherpesviridae* possess very large DNA genome (up to 235kb^1^). Herpesviruses in human encompass nine members, including HSV1/HSV2 responsible for fever blister and genital lesions, hCMV responsible for congenital malformation and a broad variety of pathologies in immunocompromised patients (graft, AIDS), Varicella Zoster Virus (VZV) that causes the childhood disease varicella (chicken pox) and establishes lifelong latency in neurons. The virus may reactivate years later and manifest as herpes zoster (shingles)^2^. Herpesviruses infection also provokes in animal severe pathologies such as Equine Herpesviruses EHV1/4 in horses that trigger rhinopathy and neurological disorder with a recent outbreak in 2021^3,4^. Pseudorabies Virus (PRV) is another veterinary herpesvirus that massively impacts swine with a potential zoonotic probability and disastrous economic consequences^5^. Few antivirals have been developed against herpesviruses. Acyclovir (ACV) is used as a treatment of HSV1 and HSV2 infections but is barely efficient on other herpesviruses. Last generation of antivirals includes ganciclovir and cidofovir with their formulated oral forms valganciclovir and brincidofovir. Ganciclovir (GCV) is a nucleoside analog that has been specifically designed to inhibit hCMV replication and is used as a curative treatment for hCMV related diseases and prophylactic treatment in case of transplantation. Cidofovir is a broad-spectrum nucleoside analog impacting replication of herpesviruses, adenoviruses and poxviruses. Due to adverse toxicity event, cidofovir use is restricted and a more bioavailable compound such as brincidofovir has been developed. Unfortunately, compassionate use of brincidofovir in three patients infected with Monkeypox virus (MPXV) recently triggered elevated liver enzymes and treatment had to be stopped^6^. Also, BCDV treatment failed in Phase III clinical trial for the management of hCMV infection in hematopoietic stem cell transplantation^7^. Letermovir (LMV) is a highly efficient inhibitor of hCMV terminase that has been approved by health authorities for the prophylaxis treatment of hCMV infection. Foscarnet (FOS), a pyrophosphate analog, is also used in the clinic in second line as a broad-spectrum antiviral perturbing phosphate binding pocket and polymerase activity^8^. Unfortunately, ganciclovir and letermovir treatment lead to the apparition of resistant strains and patients enter into therapeutic failure. There is therefore a constant need to develop new antiviral molecules.

In this study, we present the testing of the LAVR-289 prodrug, an acyclonucleoside phosphonate displaying highly efficient antiviral activities on a large collection of herpesviruses^9^. We show that LAVR-289 is active on hCMV clinical strain and gold standard resistant strains. On HSV1, LAVR-289 was able to prevent infection and protect cells from virus induced death at nanomolar concentrations. LAVR-289 also prevents HSV1 replication in reconstituted human skin model. Remarkably, LAVR-289 maintained its antiviral activity in relevant patient primary isolates, even on acyclovir and foscarnet HSV1/HSV2 resistant strains. LAVR-289 is a promising compound with remarkable activities, more potent than the current gold standard used in the clinic that paves the way for new generation broad-spectrum inhibitors of dsDNA virus infections.

## Methods

### Cells and Viruses

MRC5 (CCL-171) cells were grown in DMEM without phenol red (Sigma Aldrich) supplemented with 10 % fetal bovine serum (FBS) (Eurobio-Scientific), 1mM sodium pyruvate (S8636; Sigma Aldrich), L-Glutamine (G7513; Sigma Aldrich) and Penicillin-Streptomycin solution (P0781; Sigma Aldrich). Cells were incubated at 37°C in humidified 5% CO_2_. hCMV ANCHOR, hCMV ANCHOR GCV^R^, hCMV ANCHOR LMV^R^ were already published and produced in MRC5 cells^10^. EHV1 and EHV4 were received from ANSES. PRV was developed in collaboration with Prof Qiu laboratory and submitted elsewhere^11^. HSV1 and HSV2 strains were received from the French National Reference Center for herpesviruses, produced in MRC5 cells and titrated using high content imaging or CPE reduction assay.

### HCS imaging

For ANCHOR tagged viruses, cells were fixed with Formalin (Sigma) for 10 min at RT at the according time post-infection and washed with PBS. PBS Hoechst 33342 (1µg/mL) was then added to the cells. Cells were imaged using a Thermo CellInsight CX7 HCS microscope and results obtained using a modified compartmental spot detector algorithm. Infection rate is expressed as the percentage of GFP+ cells over the total number of cells (number of cells corresponding to the total number of cell nuclei assessed using the Hoechst 33342 signal), viral DNA replication rate is determined as the intensity and surface of the replication center. For HSV1, infected cells were detected by immunofluorescence using an antibody against ICP8 (Abcam, ab20194) and quantified using the same compartmental analysis algorithm. For HSV2 and EHV1, antiviral effect was determined using the automatic visualization and quantification of CPE using brightfield acquisition and a proprietary algorithm.

### HSV1 infection in reconstituted human skin

Foreskin fibroblasts were seeded in 24-well plates (50,000/well) and grown for 4 weeks in DMEM medium containing 10% FBS, 100 UI/ml penicillin, 100 µg/mL streptomycin and 50 µg/mL ascorbic acid (Sigma-Aldrich) to form cellular sheets. HPK (Human Primary Keratinocytes) cells were then seeded (200000 cells/well) onto fibroblast sheets in Dermalife K medium (Cellsystems GmbH). After 48 h, three sheets of fibroblasts and one sheet of HPK/fibroblasts were stacked, lifted to the air-liquid interface for epidermis differentiation and cultured in 3:1 DMEM:Ham’s F-12 medium (ThermoFisher Scientific) supplemented with 10% FBS, 0.4 µg/mL hydrocortisone (Sigma-Aldrich), 5 µg/mL bovine insulin (Sigma-Aldrich), 10^−10^ M cholera toxin (Sigma-Aldrich), 100 UI/ml penicillin, 100 µg/mL streptomycin and 50 µg/mL ascorbic acid. Media were changed three times a week. HRS (Human Reconstructed Skin) were maintained in culture for two weeks^12,13^. Skin were infected by needle injection of 10^6^ pfu of HSV1 17+ strain and treated or not with LAVR-289 at 3μM for 5 days. After fixation, an immunofluorescence against ICP8 was conducted. Image were taken on a Zeiss Axiovert A1 fluorescence microscope (40X).

## Results

### LAVR-289 inhibits hCMV replication at nM concentration and is active against GCV^R^ and LMV^R^ strains

LAVR-289 (Fig. 1A) is a new compound of the acyclonucleoside phosphonate family (4-phosphono-1-yl acyclic nucleoside), a nucleoside analogue whose osidic part is replaced by an acyclic chain. Although their structural similarity with naturally occurring nucleosides is relatively low, it has been shown that these compounds can block DNA replication by chain terminator effect or by inhibiting the activity of the viral DNA polymerase^14,15^. To determine if LAVR-289 is able to inhibit viral replication, we used this compound in an antiviral assay to determine its efficiency against hCMV infection. We used the ANCHOR tagged hCMV TB40-E strain that has already been used to detect and quantify antiviral molecules and is suitable for high content imaging^10^. ANCHOR tagged hCMV infection in permissive cells triggers the apparition of Or-GFP signal in cells so that all cells that are GFP positive are infected. Presence of the ANCH sequence in the viral genome leads to the creation of a fluorescent spot that corresponds to hCMV genome position. hCMV replication generates fluorescent spots accumulation inside the replication centers and the number and intensity of spots can be used to calculate virus replication level directly in living infected cells^10^. Infection of primary cells with hCMV ANCHOR and treating with LAVR-289 did not have any impact on the infection rate (number of GFP positive cells over total number of cells) (Fig1A, left) during the first round of infection, as expected for a compound that targets viral replication. Quantification of hCMV replication level in infected cells shows a massive effect of LAVR-289, where replication is completely abolished at concentration above 600nM with a calculated EC_50_ (Effective Concentration triggering 50% inhibition) of 40nM (Fig1A, right). Ganciclovir EC_50_ on hCMV is 2300nM^10^. Parallel cytotoxicity testing in MRC5 gave a CC_50_ (Cytotoxic Concentration triggering 50% cell death) of 25μM (*data not shown*) which corresponds to a selectivity index (CC_50_/EC_50_) of 625. High resolution imaging of cells infected with hCMV ANCHOR shows in untreated condition massive replication centers in the nucleus of infected cells (Fig1B, left). Imaging of cells infected with hCMV ANCHOR and treated with LAVR289 at 5μM confirms that the compound could not prevent hCMV ANCHOR infection because cells are still GFP positive. However, a complete loss of replication centers can be seen (Fig1B, middle). Tenfold exposure shows the presence of early replication centers containing few hCMV ANCHOR genomes, indicating that albeit cells are infected and genome was injected in the nucleus, hCMV replication did not fire (Fig1B, right). To determine if LAVR-289 could impact infection and replication of ganciclovir resistant and letermovir resistant strains, we used two strains of hCMV, hCMV ANCHOR GCV^R^ strain (mutations D301N in UL54 viral DNA polymerase catalytic subunit gene and C603W in UL97 serine/threonine kinase gene, known positions conferring Ganciclovir resistance)^16^ and hCMV ANCHOR LMV^R^ strain (mutation R369G in UL56 terminase complex subunit gene, a known position conferring Letermovir resistance)^17^. Remarkably, LAVR-289 maintains its activity in these two strains, LAVR-289 EC_50_ is 20nM on GCV^R^ strain and 50nM on LMV^R^ strain (Fig. 1C). hCMV can also cause retinitis in immunocompromised patients (AIDS) and is the first cause of blindness in the US^18^. To test if LAVR-289 can impact hCMV infection and replication in retinal cells, we used arising retinal pigment epithelial cell line (ARPE-19). Cells were infected with a high MOI of 10 and infection was carried out for 7 days, because ARPE-19 cells are known to be resistant to hCMV infection produced in fibroblast^19^. LAVR-289 was able to decrease replication level in ARPE-19 with EC_50_ of 680nM, impacting infection propagation of the virus in the cell population (Fig. S1A).

**Figure 1:**
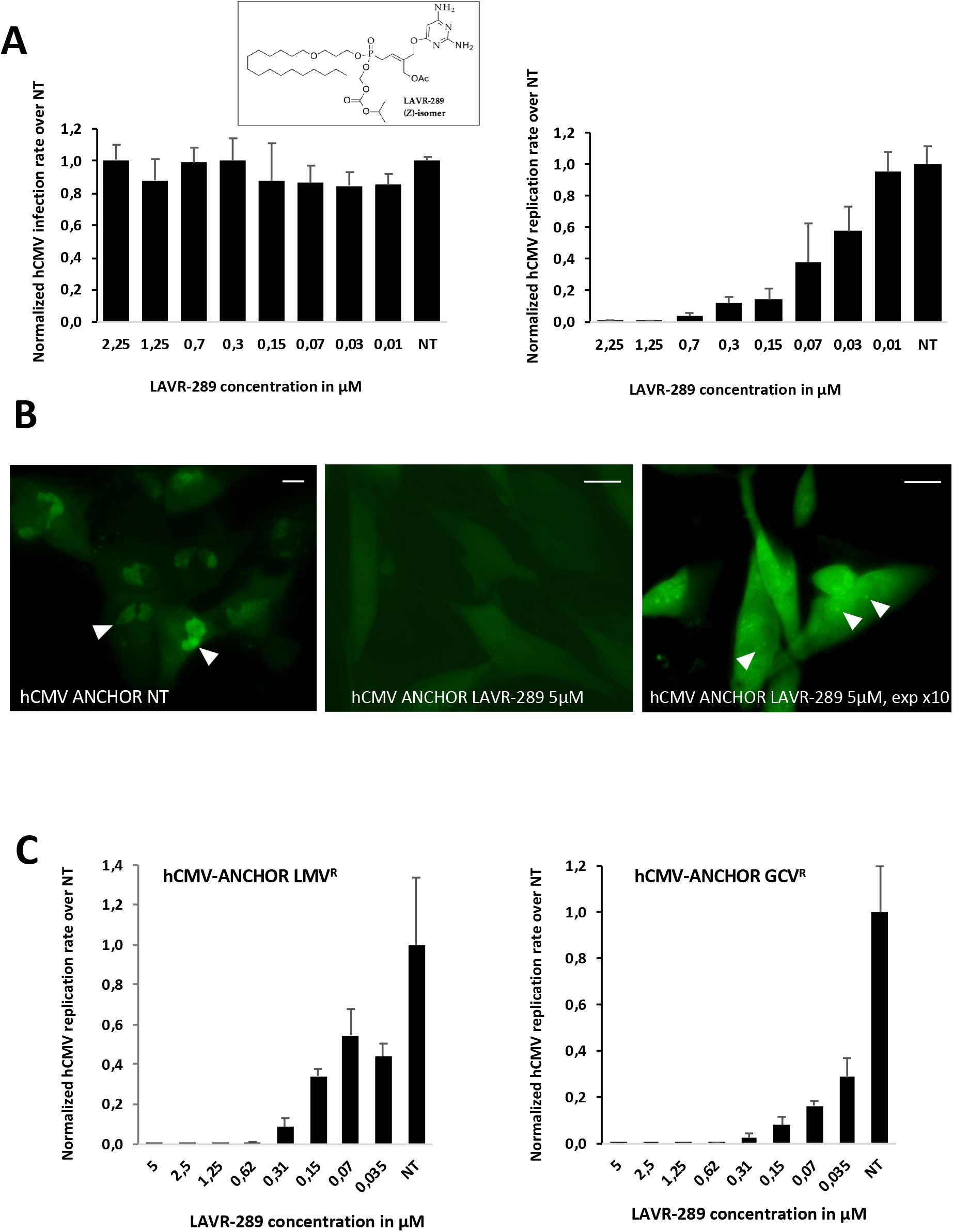
LAVR-289 inhibits hCMV replication at nanomolar concentration and interaction with existing treatments. A) MRC5 cells were infected with hCMV ANCHOR at MOI 0.5 and fixed 48h PI. Infection rate (left) and viral DNA replication level (right) were assessed for LAVR-289 treated cells at the indicated concentrations and normalized over control, untreated conditions. B) MRC5 cells were infected with hCMV ANCHOR at MOI 0.5 and imaged 72h PI. Untreated cells display massive replication centers in the cell nucleus (left, arrowheads). LAVR-289 was used at 5μM to have the terminal phenotype. LAVR-289 treated cells (middle) were processed under the same condition in terms of exposure and post-processing than the untreated control. No replication centers can be detected. Increasing exposure 10 times (right) shows that LAVR-289 treated cells are blocked at the early hCMV replication center stage (arrowheads). C) Replication rate of hCMV in MRC5 cells infected with hCMV ANCHOR LMV^R^ (left) or GCV^R^ (right) at MOI 0.5 and fixed 6d PI. The calculated IC_50_ are 50nM and 20nM for LMV^R^ and GCV^R^ strains, respectively.

### LAVR-289 inhibits propagation of clinical HSV1 and HSV2 isolates at nM concentration, is active against ACV^R^ and ACV^R^/FOS^R^ strains and prevents HSV1 progression in reconstituted human skin

To determine if LAVR-289 is able to inhibit HSV1 and HSV2 in cell culture, we used this compound in an antiviral assay to determine its efficiency on patient derived strains in collaboration with the French National Reference Center for herpesviruses. We challenged the LAVR-289 on a HSV1 clinical strain with normal sensitivity to ACV and FOS, a clinical strain exhibiting mutations in the thymidine kinase gene (C336Y) and the DNA polymerase gene (A719V) conferring ACV^R^, and a clinical strain exhibiting mutations in the thymidine kinase gene (R176Stop) and in the DNA polymerase gene (S724N) conferring both ACV^R^ and FOS^R^. MRC5 cells were plated at 12k/well, 24h post seeding cells were treated with LAVR-289 at the indicated concentrations and infected with 0,01μL (∼ 500ffu) of corresponding viruses. 48h post-infection, cells were fixed and processed for immunofluorescence against ICP8, a ssDNA binding protein required for HSV1 replication REF (Fig. 2A). Total number of cells and percentage of infected cells is assessed using high content imaging using a compartmental algorithm measuring the number of nuclei and level of ICP8 positive nuclei, respectively (Fig. 2A). Results are presented as total number of cells and percentage of infection (ICP8 positive nuclei/total nuclei) normalized over non-treated (NT) conditions. In NT conditions, cytopathic effect (CPE) reaches 100% and all cells display an ICP8 signal. Presence of LAVR-289 triggers a dose dependent decrease of the number of ICP8 positive cells, correlated with an increase in the total number of cells in the treated conditions (Fig. 2B). These results show that LAVR-289 is able to prevent HSV1 infection progression in cell culture and protect cells from virus induced death. Calculated EC_50_ on HSV1 clinical strain is 211nM. Presence of ACV^R^ mutations did not modify the calculated EC_50_ of LAVR-289, displaying 235nM EC_50_ on the A719V/C336Y ACV^R^ strain. However, mutations conferring both ACV^R^ and FOS^R^ triggered a 2.5-fold increase in LAVR-289 EC_50_ (571nM in the S724N/R176Stop ACV^R^ and FOS^R^ strain). To verify that LAVR-289 is able to inhibit HSV1 infection on more complex systems, we used reconstituted human skin. Skin was infected by needle injection of HSV1 and treated or not with LAVR-289 (at 3μM). Five days post infection, skin tissues were fixed and processed for immunofluorescence against ICP8. Whereas in NT condition HSV1 massively infected the skin (Fig. 2C, left), presence of LAVR-289 completely abolished infection (Fig. 2C, right)

**Figure 2:**
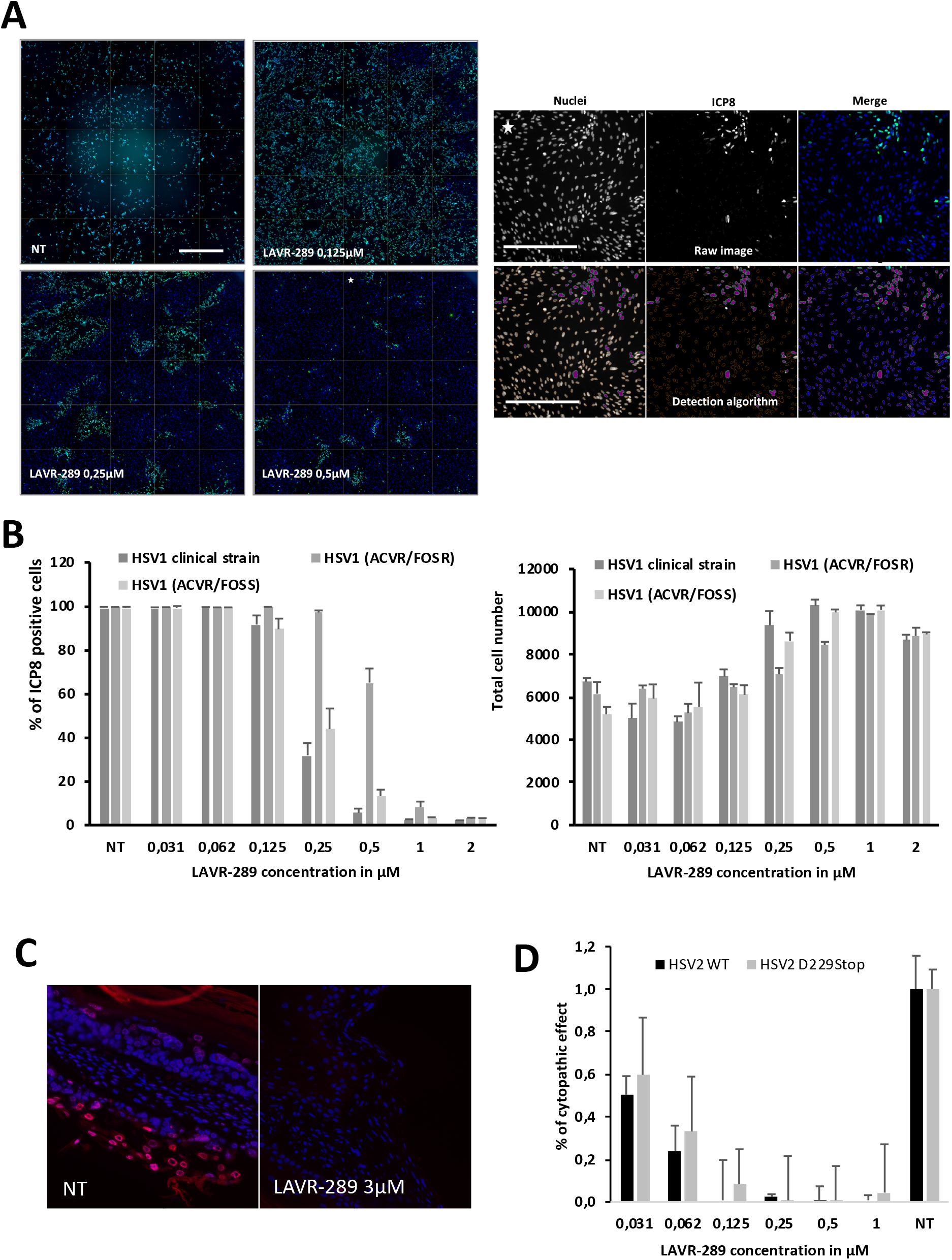
LAVR-289 inhibits HSV1 clinical strain propagation and is active on ACV^R^ and ACV^R^/FOS^R^ clinical strains, HSV1 in reconstituted human skin models and LAVR-289 inhibits HSV2 clinical strains propagation. A) Imaging of MRC5 cells infected with a clinical HSV1 strain (around 500ffu) fixed and processed for immunofluorescence against ICP8 48hPI in the presence of indicated concentrations of LAVR-289. For each condition, 25 fields reconstruction of Hoechst 33342 (blue)/ICP8 (green)s pictures are shown (bar=800μm). Example field shown in (A right panel) is indicated by a star. Example of the detection algorithm used to detect ICP8 positive cells. Raw images are presented on the left and with detection algorithm on the right. Orange object are detected as valid nuclei but negative for ICP8, green and purple object are detected positive for ICP8 (bar=400μm). B) Percentage of ICP8 positive cells (left) and total number of cells (right) according to the nature of the HSV1 virus (as indicated) and the concentration of LAVR-289 48hPI.C) in vitro reconstituted human skin samples were infected by injection with HSV1 17+ strain and left untreated or treated with LAVR-289 at 3 μM for 5 days before fixation and immunofluorescence against ICP8 (red), nuclei in blue. D) MRC5 cells were infected with HSV2 or HSV2 D229 stop (ACV^R^). 48h post infection, cytopathic effect was assessed using compartmental algorithm and normalized over NT conditions.

Same results were obtained on HSV2 or HSV2 ACV^R^ clinical samples with no detectable CPE above 250nM (Fig 2D). In a “One Health” approach, we also tested the ability of LAVR-289 to inhibit animal Herpesviruses such as EHV1(Fig. S1B), EHV4 and PRV resulting in EC_50_ of 50nM, 100nM and 400nM respectively (*data not shown*). Interestingly, the compound did not display major toxicity at high concentration (up to 10μM) nor have effect on cell permeability and mitochondrial potential in equine cells (Fig. S1C).

## Discussion and summary

Herpesvirus infection is a common infection in human population. After primary infection, virus will remain in a latent form life-long and may reactivate. HSV1 reactivation causes fever blisters, which is benign but painful. Other viruses such as hCMV can reactivate and propagate to generate a systemic infection that can be life threatening if not correctly treated either by therapeutic or prophylactic regimen. Ganciclovir and letermovir are the gold standards for hCMV treatment in clinic, but resistant strains can emerge as between 5 and 10% of patients will develop resistance and therapeutic failure in the clinic^20^. There is therefore a constant need for best in class or first in class molecules.

In this study, we have tested the efficacy of LAVR-289, a new acyclonucleoside phosphonate inhibiting virus replication by acting as a fraudulent block during viral DNA synthesis. LAVR-289 was able to inhibit a large collection of Herpesviruses from human and animal origin without toxicity. Imaging of cells infected with hCMV and treated with LAVR-289 shows that even if cells are infected, they are blocked at the early replication stage, confirming that LAVR-289 inhibits virus replication firing. LAVR-289 displays an EC_50_ of 40nM on the hCMV TB40-E clinical strain, compared to the 2.3μM of ganciclovir. LAVR-289 is therefore more than 50 times more potent than ganciclovir *in vitro*. Importantly, LAVR-289 has been designed as a pro-drug and therefore 2/3 of the mass of the compound is composed of biolabil groups that should not have any direct activities on viral DNA polymerase despite liberating the acyclic nucleoside intracellularly. The effective EC_50_ should therefore be around 13nM, making LAVR-289 170 more potent than ganciclovir to inhibit hCMV replication *in vitro*. LAVR-289 was also able to mitigate virus propagation in retinal cell line, this could be interesting to develop eye drops or intravitreal administration for the management of hCMV retinitis. Remarkably, its activity is maintained on hCMV strains resistant to ganciclovir or letermovir, offering new perspectives for the treatment of patient infected with either GCV^R^ or LMV^R^ viruses. This also opens new perspectives in the treatment of hCMV induced congenital malformation, as hCMV primary infection during pregnancy (mainly in the first trimester) can lead to fatal or severe defect in newborn children.

LAVR-289 is active on all HSV1/2 clinical strains tested so far, independently of their resistance to gold standards, albeit Foscarnet resistant strain is less sensitive to LAVR-289. Foscarnet is a pyrophosphate analog that inhibits chain elongation by competing with pyrophosphate exchange inside the polymerase. It is therefore interesting to see that phosphate metabolism in the viral DNA polymerase is also important for LAVR-289 activity on Herpesviruses, as it is suggested for poxviruses polymerases (see Marcheteau et al. same issue). LAVR-289 also inhibited HSV1 virus propagation in reconstituted human skin, and topical formulation could be designed to treat fever blisters for example. *In vivo* administration of LAVR-289 by the subcutaneous route (Marcheteau et al., same issue) or *per os* (Quentin-Froignant et al. same issue) was well tolerated in mice and hamster and did not display any behavioral changes, weight loss or any sign of distress, with normal plasmatic parameters. According to its pharmacochemical properties and design, LAVR-289 can be administered *per os* and does not need any IV injection contrary to ganciclovir. Also, in a one-health approach, LAVR-289 was tested on Herpesviruses of veterinary interest and displayed nanomolar activities on EHV1, EHV4 and PRV. This can be very interesting for the management of EHV outbreaks and to treat and protect equestrian centers. Also, treating PRV infected swine farm could help to reduce the viral burden and decrease the probability of virus propagation to surrounding farms, and therefore herd culling.

Having a compound displaying high efficacy *in vivo* with *per os* bioavailability and maintenance of activity on resistant strains will be a great asset to tackle any herpesvirus related diseases and increase the chance for full recovery of patient with herpesvirus opportunistic infection and more severe and acute herpes associated encephalitis.

## Acknowledgments

Authors thank French Defense Innovation Agency (RAPID program “Denalpovir” grant # 192906106), Region Centre Val de Loire (APR-IR FINALS) for financial support, which made this study possible. ICOA UMR CNRS 7311 receives grants from the University of Orléans and from the CNRS as well as from ANR Precyvir and FEDER (EX003677, EX011313, 2021-2027-00022860), Labex SYNORG (ANR-11-LABX-0029) and IRON (ANR-11-LABX-0018-01), and the SALSA platform for spectroscopic measurements.

## Author Contributions

Conceptualization: F.G.; Experiments: S.K.G, C.Q.F., E.M, J.F., D.B., V.G., V.R, L.A.A., Supervision: F.G.; Writing original draft: F.G. – review&editing: F.G., D.B., L.A.A. Funding acquisition: L.A.A., F.G. All authors have read and agreed to the published version of the manuscript.

## Figure legend

**Figure S1:**
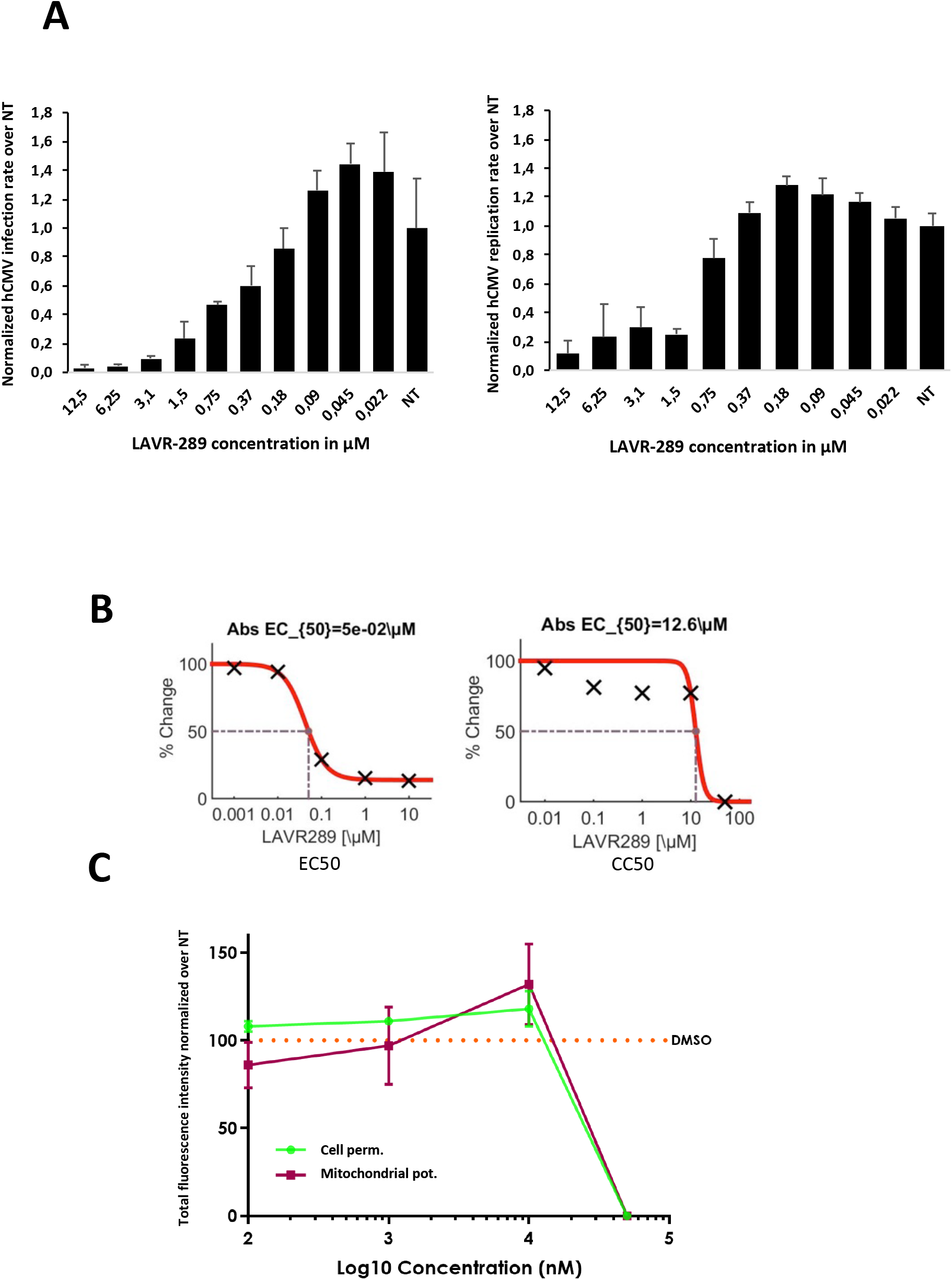
LAVR-289 inhibits hCMV in retinal cells and EHV1 in Equine dermal fibroblasts. A) ARPE-19 cells were infected with a high MOI of 10 and infection was carried out for 7 days. Infection rate (left) and viral DNA replication level (right) were assessed for LAVR-289 treated cells at the indicated concentrations and normalized over control, untreated conditions B) EC_50_ calculation at 50nM and CC_50_ at 12,6 μM gives a selectivity index of 252, on ED cells infected with EHV1 virus Calculation were done on Combenefit C) Effect of LAVR-289 on cell permeability and mitochondrial potential assessed by high content microscopy, using the cell toxicity kit from Thermo Scientific. No major effect was visualized below 10 μM.

